# A highly attenuated Vesiculovax vaccine rapidly protects nonhuman primates against lethal Marburg virus challenge

**DOI:** 10.1101/2022.01.22.477345

**Authors:** Courtney Woolsey, Robert W. Cross, Krystle N. Agans, Viktoriya Borisevich, Daniel J. Deer, Joan B. Geisbert, Cheryl Gerardi, Theresa E. Latham, Karla A. Fenton, Michael A. Egan, John H. Eldridge, Thomas W. Geisbert, Demetrius Matassov

**Affiliations:** Department of Microbiology and Immunology, University of Texas Medical Branch, Galveston, TX 77555, USA; Galveston National Laboratory, University of Texas Medical Branch, Galveston, TX 77555, USA; Department of Viral Vaccine Development, Auro Vaccines, Pearl River, New York 10965, USA; Department of Immunology, Auro Vaccines, Pearl River, New York 10965, USA.

**Keywords:** Marburg virus, filovirus, rVSV, VSV, Vesiculovax, ring vaccination, Ervebo, Angola variant

## Abstract

**Background:** Marburg virus (MARV), an Ebola-like virus, remains an eminent threat to public health as demonstrated by its high associated mortality rate (23-90%) and recent emergence in West Africa for the first time. Although a recombinant vesicular stomatitis virus (rVSV)-based vaccine (Ervebo) is licensed for Ebola virus disease (EVD), no approved countermeasures exist against MARV. Results from clinical trials indicate Ervebo prevents EVD in 97.5-100% of vaccinees 10 days onwards post-immunization.

**Methodology/Findings:** Given the rapid immunogenicity of the Ervebo platform against EVD, we tested whether a similar, but highly attenuated, rVSV-based Vesiculovax vector expressing the glycoprotein (GP) of MARV (rVSV-N4CT1-MARV-GP) could provide swift protection against Marburg virus disease (MVD). Here, groups of cynomolgus monkeys were vaccinated 7, 5, or 3 days before exposure to a lethal dose of MARV (Angola variant). All subjects (100%) immunized one week prior to challenge survived; 80% and 20% of subjects survived when vaccinated 5- and 3-days pre-exposure, respectively. Lethality was associated with higher viral load and aberrant innate immunity signaling, whereas survival correlated with development of MARV GP-specific antibodies and early expression of NK cell-, B-cell-, and cytotoxic T-cell-related transcriptional signatures.

**Conclusions/Significance:** These results emphasize the utility of Vesiculovax vaccines for MVD outbreak management. The highly attenuated nature of rVSV-N4CT1 vaccines, which are clinically safe in humans, may be preferable to vaccines based on the same platform as Ervebo (rVSV “delta G” platform), which in some trial participants induced vaccine-related adverse events in association with viral replication including arthralgia/arthritis, dermatitis, and cutaneous vasculitis.

**Author Summary:** Marburg virus (MARV) is one of the deadliest viruses known to man. One of the most effective vaccines against this pathogen uses a recombinant vesicular stomatitis virus (rVSV) platform to express MARV glycoprotein (GP) immunogen. As rVSV-based vaccines may be used as medical interventions to mitigate or prevent outbreaks of MARV, defining the time window needed to elicit protection is vital. Here, a rVSV vector expressing MARV glycoprotein (rVSV-N4CT1-MARV-GP) fully protected nonhuman primates from lethality and disease when given as soon as 1 week prior to exposure. At 5- and 3-days pre-exposure, partial protection (80% and 20% survival, respectively) was achieved. Vaccination with rVSV-N4CT1-MARV-GP appears to “jump-start” the immune system to allow sufficient time for MARV-specific adaptive responses to form. This fast-acting vaccine is based on a similar platform as Ervebo, the only FDA- and EU-approved vaccine for preventing Ebola virus infection. The rVSV-N4CT1-MARV-GP vaccine features additional attenuations in the rVSV backbone that may contribute to a more acceptable safety profile in vaccinees, as Ervebo in some recipients induced vaccine-related adverse events including rashes and joint pain.

## Introduction

The genera *Marburgvirus* and *Ebolavirus* are cousins in the family *Filoviridae* that cause a similar life-threatening hemorrhagic disease in humans and nonhuman primates (NHPs) [1]. Due to their high risk to national security and public health, viruses in both genera are classified as World Health Organization (WHO) High Priority Category A pathogens [2] and US Centers for Disease Control (CDC) Tier 1 select agents [3]. While *Ebolavirus* contains six genetically distinct species, *Marburgvirus* contains a single species: *Marburg marburgvirus* (MARV).

In 2004-2005, MARV was responsible for one of the deadliest filovirus outbreaks to date. The virus emerged in the Uige province of Angola resulting in 252 confirmed cases and 227 deaths (∼ 90% case fatality rate) [4]. Outbreaks of MARV are primarily restricted to eastern and southern Africa, which largely overlaps with the geographic distribution of its reservoir species, the Egyptian fruit bat (*Rousettus aegyptiacus*) [5]. While MARV outbreaks have so far been limited and sporadic, field studies in Uganda indicate that 2– 3% of Rousette bats are actively infected with Marburgviruses at any given time [6]. Biannual seasonal pulses contribute to a ∼ 10% increase in MARV infections in juvenile Rousette bats that coincide with spillover into human populations [6]. This high rate of infection along with the extensive seroprevalence in Rousette bats underscore the underappreciated threat that MARV poses to public health. Marburgviruses have also recently emerged in previously non-endemic regions. On August 6^th^, 2021, the first known case of Marburg virus disease (MVD) in West Africa was reported to the World Health Organization [7]. The case originated in a villager from southwestern Guinea, not far from the Sierra Leonean and Liberian borders. Prior to the outbreak, surveillance in the region revealed evidence of filoviruses circulating in nearby Sierra Leone, with active MARV infection in approximately 2.5% of Rousette bats [8]. Virus sequences obtained from Rousette bats in this area were most genetically similar to human isolates identified during the deadly MARV-Angola outbreak (2005), as well as bat isolates in Gabon and the Democratic Republic of Congo (DRC) (2006–2009). This evidence highlights the importance of pathogen surveillance in these regions and stresses the need for medical countermeasures against MVD as spillover events will likely continue to occur.

While substantial progress has been made towards the development of vaccines for one ebolavirus species, *Zaire ebolavirus* (EBOV), no licensed MVD vaccines or therapeutics are currently available. Ervebo® is the only vaccine approved by both the US Food and Drug Administration (FDA) and European Medicines Agency (https://www.ema.europa.eu/en/medicines/human/EPAR/ervebo) and is recommended by the WHO and US Advisory Committee on Immunization Practices (ACIP) for the prevention of Ebola virus disease (EVD) [9, 10]. The vaccine is comprised of a live-attenuated, recombinant vesicular stomatitis virus (rVSV) vector that expresses EBOV glycoprotein (GP) immunogen instead of its native glycoprotein (G). Administration of Ervebo to contacts (and contacts of contacts) of confirmed cases in a ring vaccination trial during the 2013-2016 West Africa and 2018-2020 DRC EBOV outbreaks prevented disease in 97.5-100% of those immunized within 10 days onwards [11, 12]. These data demonstrate the ability of Ervebo to serve as a fast-acting vaccine. To protect exposed individuals and reduce community transmission, a similar vaccination strategy could be implemented in the event of a MARV outbreak.

Preclinical studies have demonstrated the use of rVSV vectors to serve as preventative vaccines and postexposure treatments against MVD in guinea pigs and nonhuman primates (NHPs) [13–21]. NHPs serve as the most stringent animal model for testing medical countermeasures and are considered the “gold standard” for recapitulating human manifestations of MVD including coagulopathies [22]. When administered ∼ 1 month before challenge, a single dose of a rVSV vaccine expressing MARV GP fully protected NHPs against a 1000 PFU lethal challenge of various MARV variants [16, 18, 20]. Immunity with an rVSV vector was also durable with 100% protection against a lethal MARV exposure 14 months post-vaccination [23]. Moreover, all NHPs survived when the vaccine was administered 20-30 minutes post MARV (Musoke variant) exposure at the same challenge dose [15]. However, treatment with a rVSV vector at 20-30 minutes after infection failed to fully defend macaques against challenge with the Angola variant of MARV (only 25% survival) [14]. *In vitro* studies demonstrate the Angola variant enters host cells via C-type lectin receptors more efficiently than MARV-Musoke, which may contribute to its increased pathogenicity [24]. MARV-Angola also yields a more rapid disease course and a more pronounced pathology in outbred guinea pigs [25] and in NHPs [26], suggesting the Angola variant sets a high bar for MVD vaccines and therapeutics in terms of conferring protection.

While preclinical studies have demonstrated the ability of rVSV-based vaccines to rapidly combat Marburgviruses including the Angola variant [13, 14], the exact prophylactic window remains undefined. Recently, Marzi et al. reported that a “delta G” rVSV-based vaccine (rVSVΔG-MARV-GP-Angola) similar to Ervebo fully protected NHPs when vaccinated at 7 or 14 days prior to exposure [27]. Partial protection (75%) was observed when NHPs were vaccinated 3 days before exposure. Correspondingly, postexposure treatment with a similar vaccine at 20-30 minutes after infection was shown to be 89% effective in protecting NHPs against a low 50 PFU dose of MARV-Angola [13]. Although these results are promising, safety studies with the delta G Ervebo vector have shown undesirable vaccine-related adverse events in humans including prolonged and recurrent arthritis symptoms, maculopapular or vesicular dermatitis, cutaneous vasculitis, and viral shedding [28, 29]. While Ervebo was deemed safe for human use and none of these adverse events were classified as contraindications, a highly effective but less reactogenic next generation vaccine would likely be preferred for widespread immunization.

Highly attenuated rVSV-based Vesiculovax vaccines induce low reactogenicity in humans and exhibit fast-acting potential in NHPs [13, 14, 17, 30]. When NHPs were administered rVSV-N2CT1-MARV-GP or rVSV-N4CT1-MARV-GP Vesiculovax vectors 20-30 minutes after a low dose MARV-Angola exposure (50 PFU), survival was 80% and 60%, respectively [13, 14]. Therefore, Vesiculovax vaccines provide comparable protection as delta G vaccines. Unlike delta G rVSV vectors, Vesiculovax vectors express target antigen from the first position to increase immunogen expression [17]. The native G is preserved but contains a cytoplasmic tail truncation that interferes with particle maturation due to decreased interaction of G and the nucleoprotein (N) at the nucleocapsid core [31]. The rVSV-N4CT1-MARV-GP vaccine is a highly attenuated vector containing a rVSV N4 translocation (N2 vectors contain an N2 translocation). Shuffling the rVSV N from the first to the fourth position markedly diminishes the intracellular abundance of this protein by virtue of its increased distance from the 3’ transcription promoter [32]. These modifications enable robust attenuation of the vector while retaining high immunogenicity. In this study, we tested the ability of the highly attenuated rVSV-N4CT1-MARV-GP vector to serve as a rapid-acting vaccine for reactive immunization during MVD outbreaks. NHPs were vaccinated at 7, 5, or 3 days before exposure to define the minimum time needed between vaccination and challenge to elicit protective immunity against MARV-Angola. Samples were collected over the course of the study to examine NHPs for clinical signs of disease and to characterize the immune response to vaccination after challenge.

## Methods

### Generation of rVSV vaccine vectors

The rVSV-N4CT1-MARV-GP and rVSVN4CT1-HIVgag vaccine vectors used in this study were recovered from infectious clones as described previously [14, 17]. An expression cassette encoding the full-length MARV-Angola GP (accession number: DQ447653) or HIV gag protein, respectively, was cloned into a plasmid containing the full-length VSV genome. This plasmid encodes for a VSV N1 to N4 gene translocation and VSV G CT1 truncation; the MARV-Angola GP or HIV gag gene is expressed from the first genomic position to maximize GP antigen expression. Vectors were then recovered from Vero cells following electroporation with the resulting plasmids along with VSV helper plasmids. The rescued virus was plaque purified and amplified to produce virus seed stocks. Vaccine vectors were purified and concentrated for *in vivo* experiments. Before proceeding to *in vivo* studies in NHPs, the vector genomes were completely sequenced to verify the fidelity of the open reading frames (ORFs) for all genes.

### Challenge virus

The MARV-Angola seed stock originates from the serum of a fatal patient (8-month-old female; isolate 200501379) during the 2004–2005 Uige, Angola outbreak (DQ: 447653.1). The p2 challenge material was created by passaging the original isolate 200501379 twice onto Vero E6 cells (titer 1.5 × 10^7^ PFU/mL). Stocks were certified free of endotoxin (< 0.5 EU/mL) and mycoplasma contamination.

### Ethics Statement

Animal studies were conducted in compliance with the Animal Welfare Act and other federal statutes and regulations relating to animals and experiments involving animals. All experiments adhered to principles stated in the eighth edition of the “Guide for the Care and Use of Laboratory Animals” (National Research Council, 2011). The Galveston National Laboratory (GNL) where this research was conducted (UTMB) is fully accredited by the Association for the Assessment and Accreditation of Laboratory Animal Care International and has an approved OLAW Assurance (#A3314-01). Animal studies were performed in BSL-4 biocontainment at the University of Texas Medical Branch (UTMB) and the protocol was approved by the UTMB Institutional Biosafety Committee.

### Animal challenge

Eighteen adult (9 females and 9 males) cynomolgus macaques (*Macaca fascicularis*) of Chinese origin (PreLabs, Worldwide Primates) ranging in age from 3 to 8 years and weighing 2.86 to 7.60 kg were used for three separate studies at the GNL. Macaques were immunized with a single 10 million PFU intramuscular (i.m.) injection of rVSV-N4CT1- MARV-GP at 7 (N=5), 5 (N=5), or 3 (N=5) days prior to MARV exposure. Three animals were immunized with an identical dose of rVSVN4CT1-HIVgag at each respective time point to serve as non-specific controls. The inoculation was equally distributed between the left and right quadriceps. All macaques were challenged i.m. in the left quadriceps with a uniformly lethal 1000 PFU target dose of MARV-Angola (actual doses were 1475, 1475, and 1300 PFU, respectively). An internal scoring protocol was implemented to track disease progression in challenged animals. Animals were checked at least twice daily for scoring criteria such as posture/activity level, appetite, behavior, respiration, and the presence of hemorrhagic manifestations. Subjects that reached a clinical score ≥ 9 were promptly euthanized with a pentobarbital solution.

### Blood collection

Blood was collected by venipuncture into EDTA and serum tubes pre-challenge and 3, 6, 10, 14, 21, and 28 DPI, or terminally. An aliquot of EDTA-treated whole blood (100 μl) was diluted with 600 μl of AVL inactivation buffer (Qiagen, Hilden, Germany), and RNA was extracted using a Viral RNA mini-kit (Qiagen) according to the manufacturer’s instructions. To isolate plasma and serum, tubes were spun at 2500 rpm for 10 minutes at 4°C. EDTA plasma and serum were stored at -80°C for analysis.

### Hematology and clinical chemistry

Total white blood cell counts, white blood cell differentials, red blood cell counts, platelet counts, hematocrit values, mean cell volumes, mean corpuscular volumes, total hemoglobin concentrations, and mean corpuscular hemoglobin concentrations were analyzed from blood collected in tubes containing EDTA using a laser based hematologic analyzer (VetScan HM5). Serum samples were tested for concentrations of albumin, amylase, alanine aminotransferase (ALT), alkaline phosphatase (ALP), gamma-glutamyltransferase (GGT), aspartate aminotransferase (AST), glucose, total protein, cholesterol, total bilirubin (TBIL), creatine (CRE), blood urea nitrogen (BUN), and C-reactive protein (CRP) by using a Piccolo point-of-care analyzer and Biochemistry Panel Plus analyzer discs (Abaxis).

### Viral Load Determination

One-Step Probe RT-qPCR kits (Qiagen) and CFX96 system/software (BioRad) were used to determine viral copies in samples. To detect MARV RNA, we targeted the MARV NP gene with primer pairs and a 6-carboxyfluorescein (6FAM) – 5 ′ - CCCATAAGGTCACCCTCTT-3′– 6 carboxytetramethylrhodamine (TAMRA) probe. Thermocycler run settings were 50 °C for 10 min; 95 °C for 10 s; and 40 cycles of 95 °C for 10 s plus 59 °C for 30 s. Integrated DNA Technologies synthesized all primers and Life Technologies customized the probes. Representative MARV genomes were calculated using a genome equivalent standard. The limit of detection for this assay is 1000 copies/ml.

Infectious MARV loads were determined using a standard plaque assay. Briefly, increasing 10-fold dilutions of plasma samples were adsorbed to Vero E6 monolayers (ATCC Cat: CRL-1586) in duplicate wells (200 µl), overlaid with 0.8% agarose/2x EMEM, and incubated for six days at 37 °C in 5% CO2. Neutral red stain was added, and plaques were counted after a 24- to 48-hour incubation. The limit of detection for this assay is 25 PFU/ml.

### NanoString sample preparation

Targeted transcriptomics was performed on blood samples from macaques as previously described [33]. NHPV2_Immunology reporter and capture probesets (NanoString Technologies) were hybridized with 5 µl of each RNA sample for ∼24 hours at 65°C. The RNA:probeset complexes were then loaded onto an nCounter microfluidics cartridge and assayed using a NanoString nCounter® SPRINT Profiler. Samples with an image binding density greater than 2.0 were re-analyzed with 2 µl of RNA to meet quality control criteria.

### Transcriptional analysis

Briefly, nCounter® .RCC files were imported into NanoString nSolver™ 4.0 software. To compensate for varying RNA inputs, an array of housekeeping genes and spiked-in positive and negative controls were used to normalize the raw read counts. The data was analyzed with NanoString nSolver™ Advanced Analysis 2.0 package to generate principal component (PC) figures, volcano plots, and cell-type trend plots. Human annotations were added for each respective mRNA to perform immune cell profiling within nSolver™.

Normalized data (fold-change- and p-values) were exported as a .CSV file and imported into GraphPad Prism version 9.3.1 to produce transcript heatmaps. To identify the functional annotation of individual transcripts, we interrogated the GeneCard database (https://www.genecards.org) [34]. For pathway analysis, functional enrichment of normalized counts was performed at the time of challenge, and early, mid, and late disease with Ingenuity Pathway Analysis (Qiagen). Z-scores were imported into GraphPad Prism version 9.3.1 to produce the canonical signaling heatmap.

### Anti-MARV GP IgM and IgG ELISA

MARV GP-specific IgM and IgG antibodies were quantified by ELISA on sera collected at the indicated blood collection days. Immunosorbent MaxiSorp 96-well plates were coated overnight with 15 ng/well (0.15mL) of recombinant MARV GPΔTM (ΔTM: transmembrane region absent; Integrated Biotherapeutics, Gaithersburg, MD) in a sodium carbonate/bicarbonate solution (pH 9.6). Antigen-adsorbed wells were subsequently blocked with 2% bovine serum antigen (BSA) in 1 x PBS for at least two hours. Sera were initially diluted 1:100 and then two-fold through 1:12800 in ELISA diluent (2% BSA in 1× PBS, and 0.2% Tween-20). After a one-hour incubation, cells were washed four times with wash buffer (1 x PBS with 0.2% Tween-20) and incubated for an hour with a dilution of horseradish peroxidase (HRP)-conjugated anti-rhesus IgM (1:2500) or IgG antibody (1:5000) (Fitzgerald Industries International, Acton, MA). SigmaFast O-phenylenediamine (OPD) substrate (Sigma; P9187) was added to the wells after four additional washes to develop the colorimetric reaction. The reaction was stopped with 3M sulfuric acid ∼5 minutes after OPD addition and absorbance values were measured at a wavelength of 492nm on a Cytek Cytation 5. Absorbance values were determined by subtracting uncoated from antigen-coated wells at the corresponding serum dilution. End-point titers were defined as the reciprocal of the last adjusted serum dilution with a value ≥ 0.20.

### Statistical analysis

Statistical analysis of viral load was carried out in GraphPad Prism version 9.3.1 (GraphPad, Software, Inc., La Jolla, CA) using a mixed-effects model with Geisser-Greenhouse correction and a Dunnett’s multiple comparisons test. A multiple hypothesis Benjamini-Hochberg false discovery rate (FDR) corrected p-value less than 0.05 was deemed significant for transcriptional analyses, unless otherwise stated. A Pearson correlation coefficient was employed to measure linear correlation between individual subject viremia levels and expression of specific DE transcripts.

## Results

To define the prophylactic window afforded by Vesiculovax vaccination against MVD, we immunized 15 cynomolgus macaques with a single 10 million PFU i.m. dose of rVSVN4CT1-MARV-GP at 7 (N=5), 5 (N=5), or 3 (N=5) days before challenge (**Fig 1A**). Three additional subjects were inoculated with an irrelevant rVSVN4CT1-HIV-gag1 vaccine at 7 (N=1), 5 (N=1), or 3 (N=1) days prior to MARV exposure to control for non-specific effects of the vector. All macaques were i.m. challenged with a uniformly lethal target dose of 1000 PFU of MARV-Angola and observed daily for signs of illness up to the 28 days post-infection (DPI) study endpoint.

**Fig 1.**
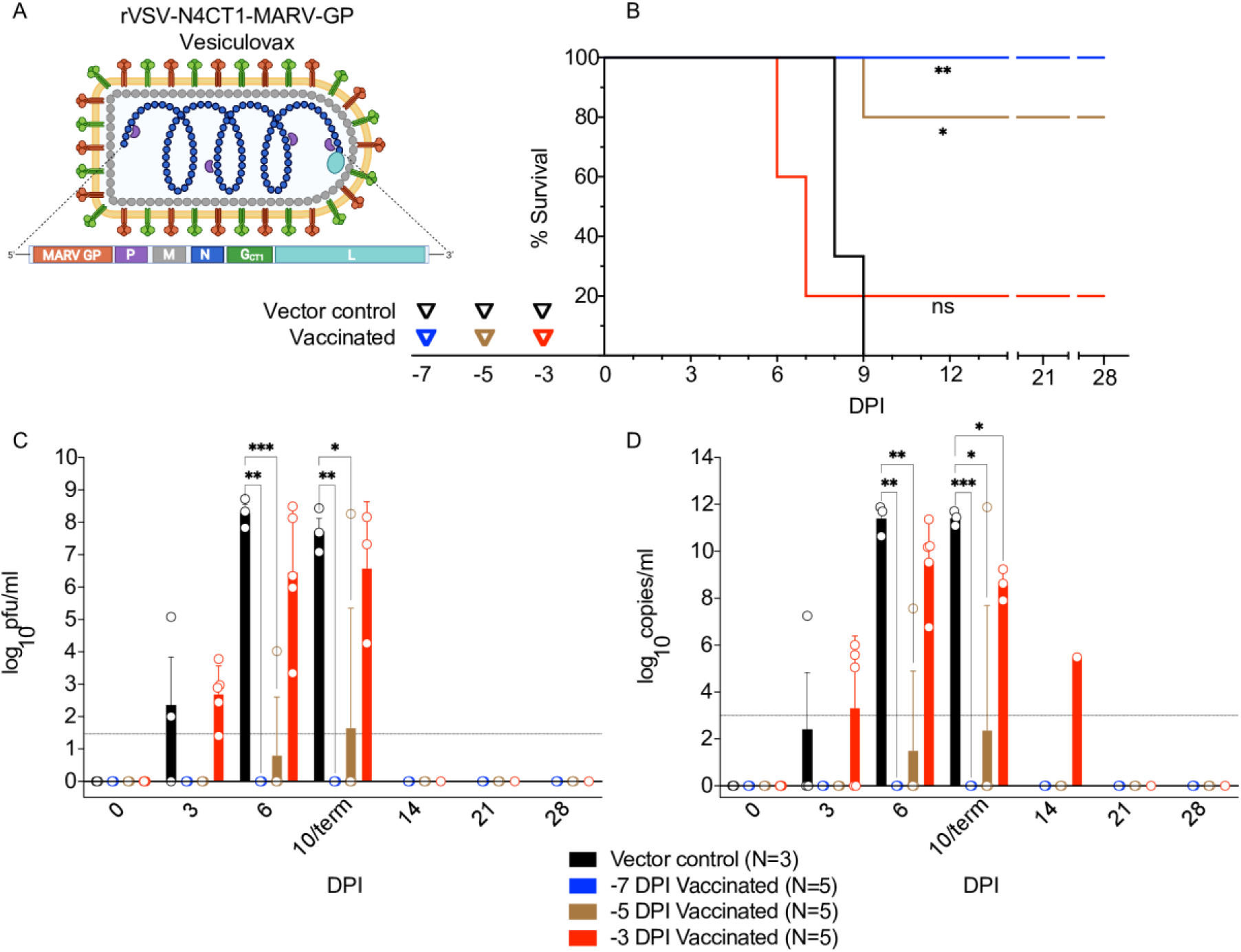
Study Design and comparison of viral loads in vaccinated macaques exposed to MARV-Angola. **(A)** rVSV-N4CT1-MARV-GP vector design and genome organization. The Vesiculovax vaccine encodes a truncated form (CT1) of the native VSV G (green box); the N gene was translocated from the 1^st^ position to the 4th position (N4) of the genome (blue box). The MARV GP glycoprotein is expressed from the 1^st^ position to increase immunogen expression (orange box). (**B**) Survival curves of vaccinated cohorts immunized with rVSV- N4CT1-MARV-GP at -7 DPI (blue; n=5), -5 DPI (brown; n=5), or -3DPI (red; n=5). A single vector control subject was vaccinated at each respective time point (black; n=3). A log-rank test was used to determine statistical significance. Triangles on the x-axis indicate time of vaccination for each respective cohort. (**C**) Plasma viremia was measured by standard plaque assay at the denoted time points and reported as log10 PFU/ml. The limit of detection for this assay is 25 PFU/ml (indicated by a dotted horizontal line). (**D**) Viral loads were measured by RT-qPCR in whole blood and reported as log10 copies/ml at the denoted time points. The limit of detection for this assay is 1000 copies/ml (indicated by a dotted horizontal line). For (**C**) and (**D**), the average titer ± SEM is shown for each group at the denoted time points. Statistical significance was determined using a mixed-effects model with Geisser-Greenhouse correction and Dunnett’s multiple comparisons test. Not significant (ns); p < 0.0332 (*); p < 0.0021 (**); p < 0.0002 (***). Abbreviations: rVSV, recombinant Vesicular stomatitis virus; MARV, Marburg virus; GP, MARV glycoprotein; VSV, Vesicular stomatitis virus; G, VSV glycoprotein; N, VSV nucleoprotein; P, VSV phosphoprotein; M, VSV matrix protein; L, VSV polymerase; DPI, days post vaccination; PFU, plaque-forming units.

Vaccination with an irrelevant or antigen-specific vector did not appear to delay the onset of disease (**Fig 1B**). The time-to-death (TTD) was 8-9 DPI for the vector controls and 6-9 DPI for fatal subjects that were specifically vaccinated (chi square test p = 0.2229). This window is in line with the typical TTD for this animal model (6-9 DPI, mean of 7.3 days) [17, 20, 35]. As expected, animals immunized earlier before challenge had a higher rate of survival. Survival rates of groups immunized with rVSVN4CT1-MARV-GP were significantly different than the vector control group with 100% (log-rank test, p = 0.0046) and 80% (p = 0.0153) efficacy for -7 DPI and -5 DPI groups, respectively. No statistical difference (p = 0.5110) was noted for the -3 DPI vaccination group, although a sole subject (20%) survived.

Regardless of the rVSV vaccine vector administered, all fatal cases presented with typical MVD clinical signs such as fever, anorexia, dyspnea, macular rash, and/or depression (**S1-S3 Tables**). Specifically vaccinated survivors remained healthy and did not display clinical signs of disease other than anorexia at 5 DPI in one subject in the -7 group (Survivor 4) (**S1 Table**) and transient anorexia and a mild petechial rash in the sole survivor (Survivor 10) in the -3 group (**S3 Table**). However, all survivors exhibited various hematological changes over the course of the study. Postmortem gross examination of fatal cases in both specifically and non-specifically vaccinated macaques revealed lesions consistent with MVD including subcutaneous hemorrhage; necrotizing hepatitis (characterized as hepatic pallor with reticulation); splenomegaly; lymphadenitis; and hemorrhagic interstitial pneumonia (characterized as failure to completely collapse and multifocal reddening of the lungs) (**data not shown**). No significant lesions were detected in examined tissues of vaccinated survivors at the study endpoint.

As anticipated, serum levels of liver enzymes and kidney function products indicative of organ damage including alanine aminotransferase (ALT), alkaline phosphatase (ALP), aspartate aminotransferase (AST), gamma-glutamyltransferase (GGT), blood urea nitrogen (BUN), and creatinine (CRE) were elevated in fatal cases (**S1- S3 Tables**). These changes were also noted in Survivor 10 of the -3 group. Lethality also corresponded with elevated CRP along with lymphopenia, thrombocytopenia, and neutrophilia.

Survival correlated with lower viral load (**Fig 1C and 1D and S1-3 Tables**). Viral titers were assessed in each cohort by performing RT-qPCR amplification of viral RNA (vRNA) and conventional plaque assays. Remarkably, neither infectious MARV nor vRNA was detected in survivors in the -7 and -5 groups (**S1 and S2 Tables**), whereas the single specifically vaccinated animal that succumbed (Fatal 1) in the -5 cohort had a viral titer of 4.02 LOG10 PFU/ml (7.56 LOG10 copies/ml) at 6 DPI. In comparison, viral loads in vector controls were 3-5 logs higher at the same time point. Similarly, a lower level of viremia was detected in the single survivor (Survivor 10) of the -3 cohort (Survivor 10) (**S3 Table**). Viral titers were comparable between fatal animals and controls at end-stage disease (∼ 6-8 LOG10 PFU/ml (∼ 9-11 LOG10 copies/ml)).

To characterize the immune response to Vesiculovax immunization, we performed targeted transcriptomics on whole blood RNA from MARV-exposed vaccinated AGMs. Spatial visualization of the dataset via principal component analyses (PCA) indicated RNA samples clustered independently of time of vaccination (group), but dimensional separation was observed for disposition (fatal, survivor) and DPI (0, 3, 6, 10/terminal) covariates (**Fig 2A**). Minimal expression changes were detected in samples from surviving and non-surviving subjects on the day of challenge. Thereafter, fatal samples exhibited timepoint- distinct clustering irrespective of whether they were derived from specifically or non-specifically vaccinated subjects, denoting similar transcriptional profiles among these animals. Most survivor samples displayed minimal spatial variation at each DPI regardless of the time of vaccination, suggesting overall expression changes in surviving subjects were modest.

**Fig 2.**
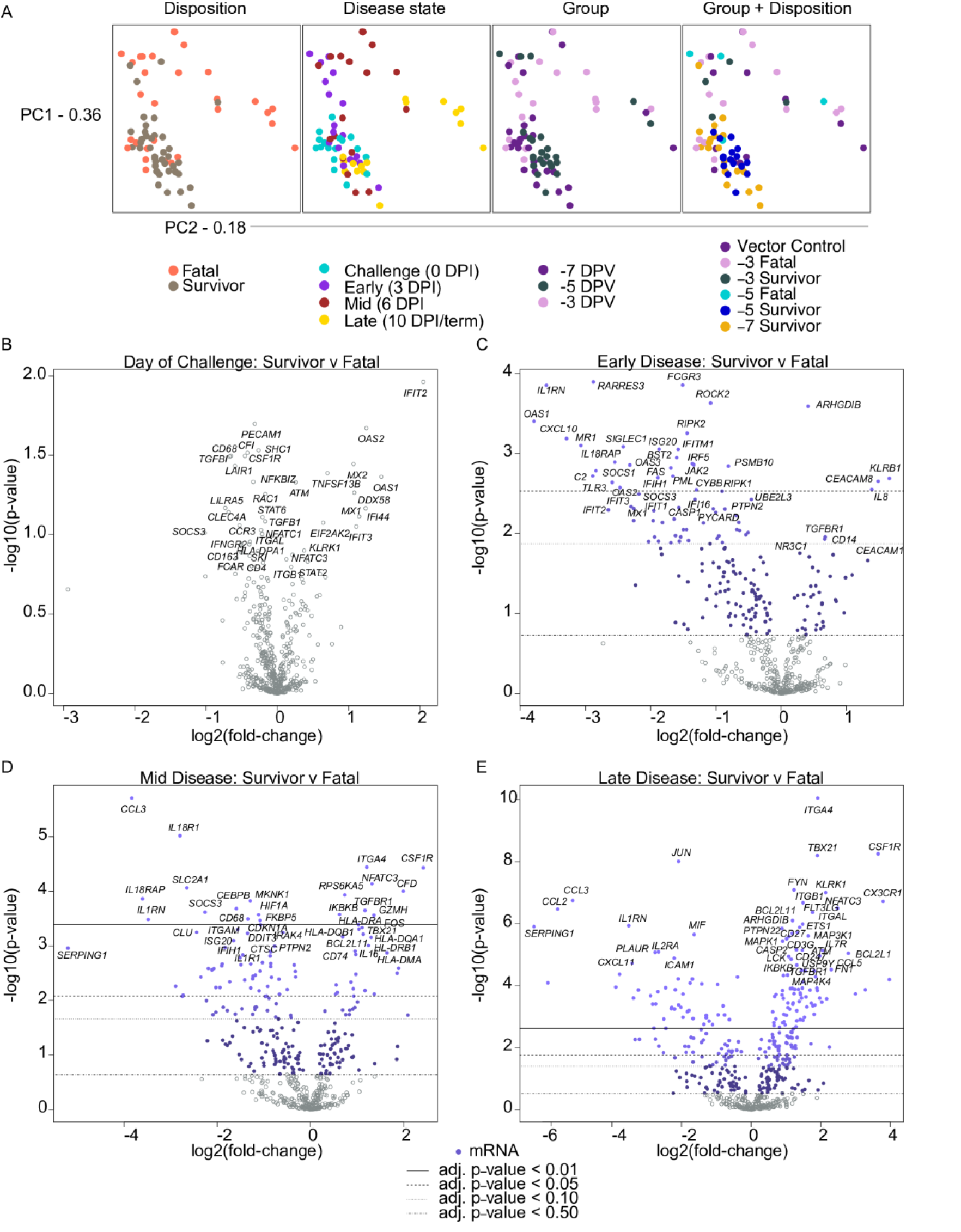
Principal component analysis and volcano plots depicting transcriptional changes in vaccinated macaques exposed to MARV-Angola. **(A)** Shown are principal component (PC) analyses of all normalized transcripts on the day of challenge (0 DPI), and at early (3 DPI), mid (6 DPI), and late disease (10 DPI, or the terminal time point in fatal cases). Individual samples were filtered by disposition (fatal, survivor), disease state (0, 3, 6, 10/term DPI), group (day of vaccination; -7, -5, or -3 DPI), and group plus disposition (vector control (n=3), -3 fatal (n=4), -3 survivor (n=1), -5 fatal (n=1), -5 survivor (n=4), -7 survivor (n=5)). **(B)** Volcano plots displaying -log10(p-values) and log2 fold changes for each mRNA target on the day of challenge (0 DPI), and at early (3 DPI), mid (6 DPI), and late disease (10 DPI, or the terminal time point in fatal cases). Horizontal lines within the plot indicate adjusted p-value thresholds. Targets highlighted in blue indicate those differentially expressed in the survivor versus fatal group irrespective of time of vaccination or vector administered with a multiple hypothesis Benjamini-Hochberg false discovery rate (FDR) corrected p-value less than 0.05. Abbreviations: PC1 (principal component 1); PC2 (principal component 2); DPI, days post infection.

To identify transcriptional signatures of protection, we performed differential expression analysis of survivor and fatal samples at each DPI. Given the lack of significant variation among group (day of vaccination) or vaccine vector administered in fatal cases, we excluded these factors for this analysis. Thus, the survivor dataset included samples from specifically vaccinated survivors at each DPI without respect to time of vaccination, whereas the fatal dataset included samples from both specifically and non-specifically non-surviving subjects at each DPI without respect to time of vaccination.

Overall, we identified 76, 70, and 168 differentially expressed (DE) transcripts (Benjamini-Hochberg (BH)-adjusted p-value < 0.05) in survivor versus fatal samples at early (3 DPI), mid (6 DPI), and late (10 DPI/terminal timepoint) disease, respectively (**S1 Data**). On the day of challenge (0 DPI), survivors tended to express higher levels of transcripts associated with interferon signaling (e.g., *IFIT2*, *OAS1*, *OAS2*, *MX1*, *MX2*, and *IFI44*) and retinoic acid-inducible gene I (RIG-I; *DDX58*) signal transduction that is involved in cytosolic detection of double-stranded viral RNA (**Fig 2B**). These findings suggest earlier activation of innate immunity in these subjects, although no statistically significant difference was found between survivor and fatal samples at this timepoint. At early disease (3 DPI), survivors expressed a higher abundance of natural killer (NK) cell- and cytolytic T lymphocyte (CTL)-affiliated transcripts (*KLRB1*, *KLRD1*, *GZMK*) as well as molecules involved in neutrophil chemotaxis and adhesion (*IL-8* and *CEACAM8*). Conversely, lower expression of inflammatory markers including *IL1RN* (encodes interleukin 1 receptor antagonist), *OAS1* (encodes 2’-5’-oligoadenylate synthetase 1), *CXCL10* (encodes C-X-C motif chemokine ligand 10, or IP-10), *MR1* (encodes major histocompatibility complex, class I-related), *RARRES3* (encodes IL-1 receptor antagonist), *C2* (encodes complement component C2), *IL18RAP* (encodes interleukin-18 receptor accessory protein) was evident in survivors (**Fig 2C**). At mid disease (6 DPI), survivors expressed more adaptive immunity-related transcripts (*ITGA4*, *NFATC3*, *TGFBR1*), antigen presentation-related molecules (*HLA-DQB1*, *HLA-DRA*, *HLA-DQA1*, *HLA-DRB1*, *HLA-DMA*), and NK cell-associated transcripts (*GZMH*). Also detected in survivors was increased expression of *CSF1R* (encodes a cytokine that controls the production, differentiation, and function of macrophages) and *TBX21* (the master transcriptional regulator for T helper 1 (Th1) cells). Decreased expression of molecules in survivors included those involved in complement regulation (*SERPING1*), pro-inflammatory signaling (*IL18RAP*, *IL18R1*, *IL1RN*), metabolism (*SLC2A1*), or recruitment and activation of monocytes/granulocytes (*CCL3*, *CD68*) (**Fig 2D**). At late disease, again higher expression of *CSF1R*, *TBX21*, and markers of NK cell and T cell activation (*CX3CR1*, *ITGA4*, *KLRK1*, *NFATC3*, *FYN*, *CD27*, *CD3G*) was noted in survivors, with lower expression of *SERPING1*, *CCL3*, *IL1RN* (**Fig 2E**).

For a more granular assessment, we examined DE transcripts for each group and disposition with respect to a pre-challenge baseline. Samples from specifically vaccinated survivors at -3 DPI and specifically vaccinated fatal subjects at -5 DPI were excluded from this analysis since these cohorts only consisted of a single subject; however, a cursory examination indicated specifically vaccinated survivor versus specifically vaccinated fatal subjects tended to express higher levels of transcripts mapping to the adaptive immunity gene set when vaccinated at 5 versus 3 days prior to challenge (**S1 Fig**). The topmost upregulated and downregulated DE transcripts in subjects vaccinated with rVSV-N4CT1- MARV-GP at -7 DPI is depicted in Fig. 3. Notably, a similar transcriptional landscape was observed in survivors immunized at -5 or -7 DPI regardless of DPI in line with our PCA results (**Fig 3A**). Vector controls and non-surviving AGMs vaccinated at -3 DPI also exhibited similar expression profiles. Abundant expression of cytoplasmic granules such as granzymes (*GZMA*, *GZMB*, *GZMK*) and perforin (*PRF1*) was apparent in survivors at all time points, suggesting early and sustained activation of NK- and cytotoxic T-cells, whereas granzyme and perforin expression in fatal subjects was not detected until mid or late disease. Th1-associated Tbet (*TBX21*) was also upregulated in the Survivor -5 and Survivor -7 datasets, whereas this transcription factor was downregulated at all time points in the Fatal -3 cohort and at mid- and late disease in the vector control group. In contrast, repressed transcripts in specifically vaccinated survivors at late disease were implicated in monocyte recruitment (*CCR1*); pattern recognition receptor signaling (*TLR5*, *MYD88*); and inflammation (*TNFAIP6*, *NFKB1A*) (**Fig 3B**). Thus, survivor transcriptional signatures correlated with predicted Th1 differentiation and activation of NK cell and adaptive responses, whereas lethality correlated with sustained innate immunity signaling and inflammation.

**Fig 3.**
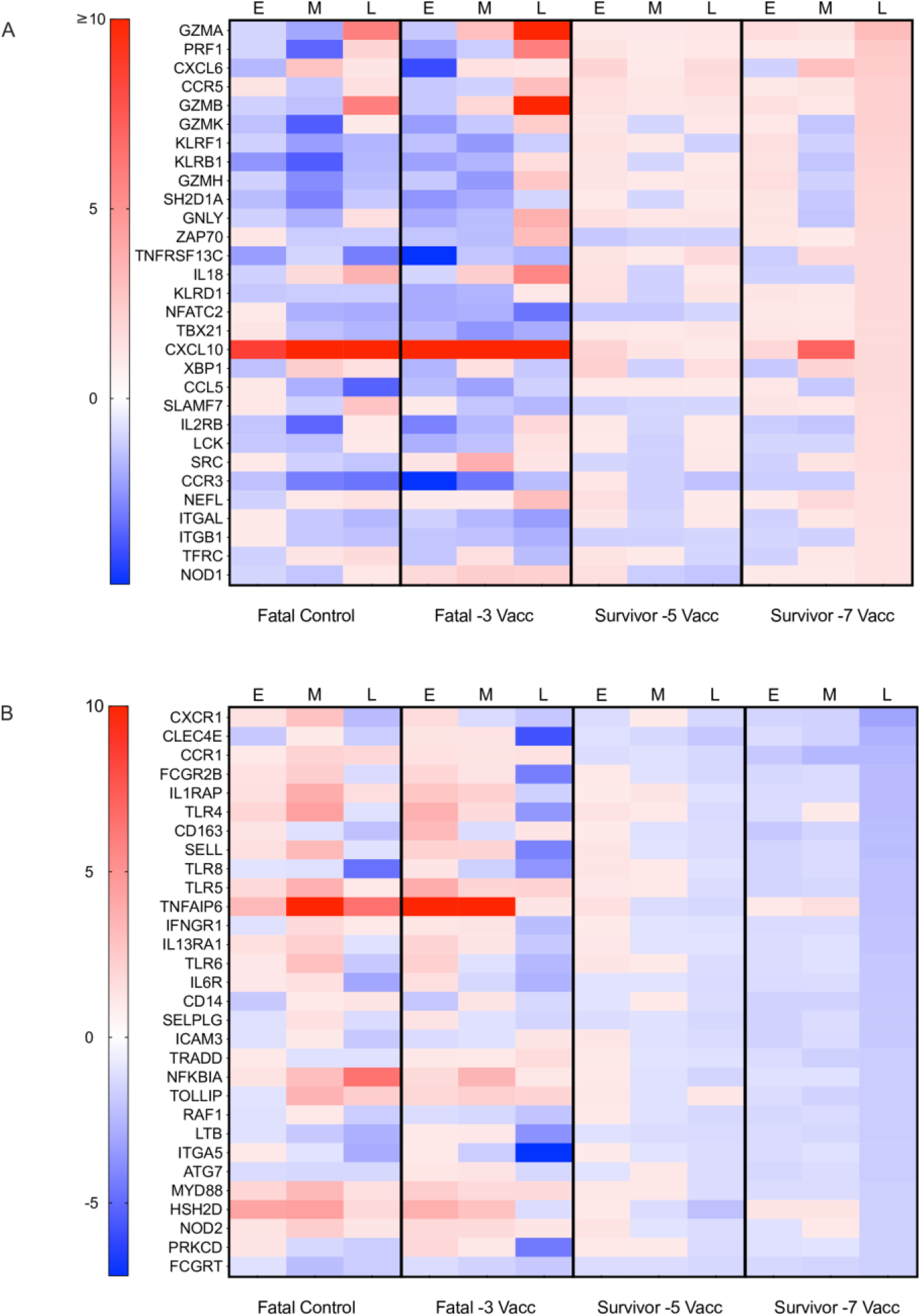
Heatmaps depicting the most differentially expressed transcripts in Vesiculovax-vaccinated survivor samples. The topmost upregulated (**A**) and downregulated (**B**) transcripts in vaccinated survivors immunized -7 DPI at late disease. Shown are expression changes in survivor cohorts (survivor -7 vacc (n=5); survivor -5 vacc (n=4)) and fatal groups (fatal vector control (n=3); fatal -3 vacc (n=4)) at each disease state (0, 3, 6, 10 DPI, or the terminal time point in fatal subjects) with respect to a pre-vaccination baseline. A Benjamini-Hochberg false discovery rate (FDR) corrected p-value less than 0.05 was deemed significant. Red indicates high expression; blue indicates low expression; white indicates no change in expression. Abbreviations: Vacc, vaccinated; E, early disease (3 DPI); M, mid disease (6 DPI); L, late disease (10 DPI or terminal timepoint).

Fatal outcome was associated with dramatic transcription of genes previously correlated with filovirus disease lethality. These transcripts included pro-inflammatory IP- 10 (*CXCL10*; up to a 65- and 55-fold-increase in vector controls and fatal subjects, respectively) [36, 37]; neutrophil-associated calgranulin A (*S100A8*; up to an 18- and 20- fold-increase in vector controls and fatal subjects, respectively) and calgranulin B ( *S100A9*; up to a 16- and 14-fold-increase in vector controls and fatal subjects, respectively) [33]; and molecules involved in immune exhaustion and anergy (*LAG3*, *CTLA4*) [38] (**S2A Fig**). Positive correlations were found between MARV viremia levels and *S100A8* (Pearson, P < 0.0001), *S100A9* (Pearson, P = 0.0001), *LAG3* (Pearson, P = 0.0021), and *CTLA4* (Pearson, P = 0.0005) counts (**S3 Fig**). Markedly diminished expression of mRNAs encoding complement factor D (*CFD*; up to a 32- and 38-fold-decrease) was observed in vector controls and fatal subjects, respectively (**S2B Fig**).

To capture shifts in circulating cell populations associated with survival, we conducted nSolver-based immune cell type profiling at 0, 3, 6, and 10 DPI (or the terminal time point in euthanized subjects). In agreement with our differential expression results, survival was associated with higher frequencies of cytotoxic cells, Th1 cells, T cells, and B cells (**Fig 4**). Lower predicted frequencies of neutrophils and macrophages were detected in fatal samples at all time points, confirming our hematology results. Contradicting our DE analysis, decreased frequencies of CD56^dim^ NK subsets were found in vaccinated survivors at certain time points. This discrepancy could reflect species-specific immunogenetic differences that exist between human and macaques [39, 40] as the nSolver profiling algorithm relies on human NK cell annotations for cell type profiling.

**Fig 4.**
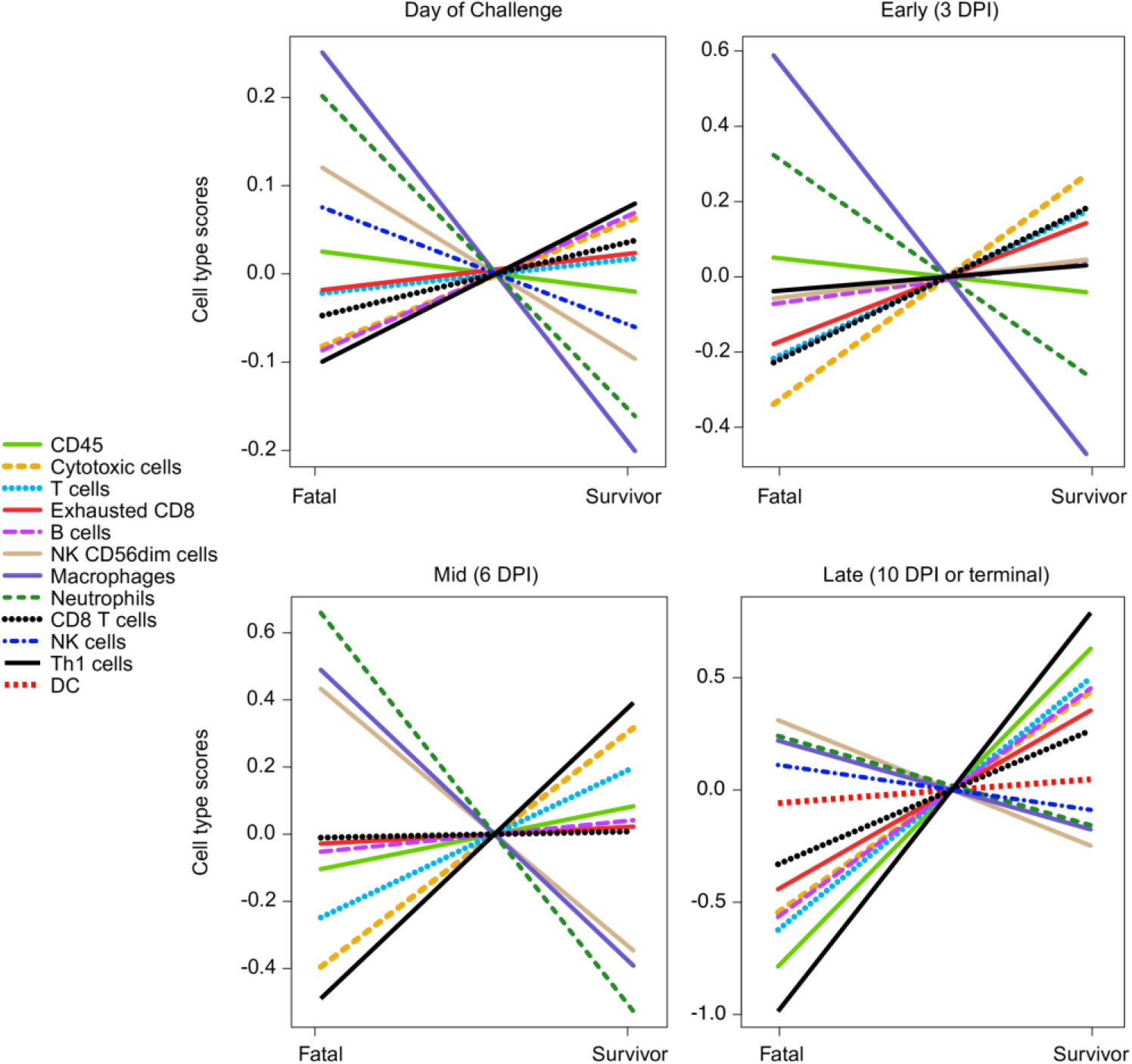
Immune cell type profiling of Vesiculovax-vaccinated survivor and fatal samples. Respective cell-type quantities for vaccinated survivor (n=10) and fatal (n=8) cohorts on the day of challenge (0 DPI), and early (3 DPI), mid (6 DPI), and late disease (10 DPI, or the terminal time point in fatal cases). A Benjamini-Hochberg false discovery rate (FDR) corrected p-value less than 0.05 was deemed significant for immune cell type profiling of transcripts. DPI, days post infection.

Next, enrichment of DE transcripts was executed to unravel canonical signaling pathways associated with rVSV-N4CT1-MARV-GP-elicited protection. At mid and late disease, the top upregulated pathways in survivor versus fatal cases included “PKC-theta signaling in T lymphocytes”, “calcium-induced T lymphocyte apoptosis”, “iCOS-iCOSL signaling in T helper cells”, “CD28 signaling in T-cell helper cells”, and “CD27 signaling in lymphocytes” (**Fig 5**). Thus, positive z-scores in survivors were associated with T cell activation and differentiation of antigen-specific T-cells. Downregulated pathways in survivors at these timepoints included those involved in innate immunity signaling (“role of pattern recognition receptors in recognition of bacteria and viruses”, “interferon signaling”, “activation of IRF by cytosolic pattern recognition receptors”) or immune dysregulation (“systemic lupus erythematous in B cell signaling pathway”, “PD-1, PDL-1 cancer immunotherapy”).

**Fig 5.**
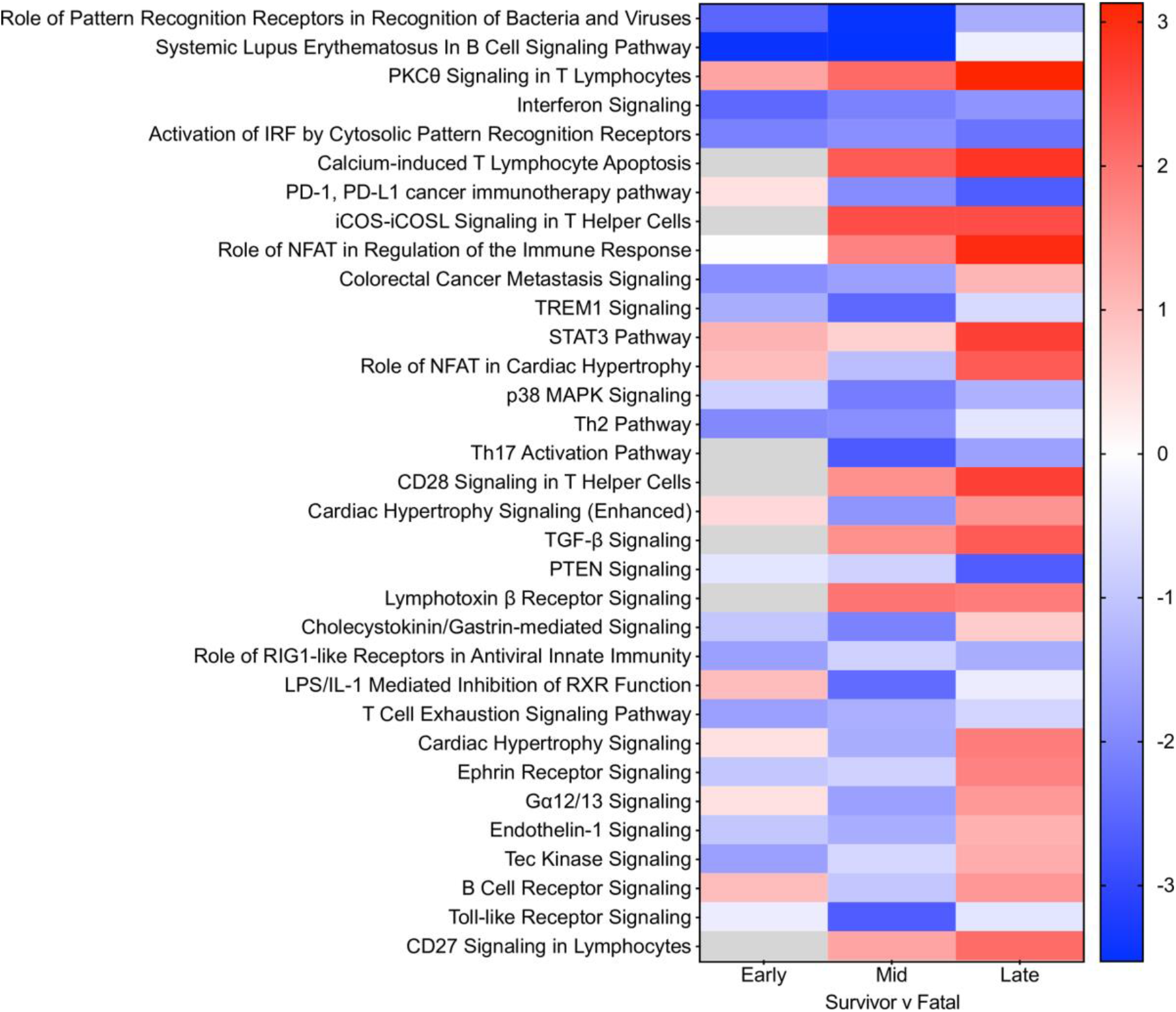
Differentially expressed canonical signaling pathways in Vesiculovax- vaccinated macaques. Heatmap of the most significantly upregulated and downregulated canonical pathways in survivor versus fatal subjects based on functional enrichment of differentially expressed transcripts (Benjamini-Hochberg false discovery rate (FDR) corrected p-value less than 0.05) in vaccinated survivor (n=10) versus fatal (n=8) samples. Cohorts were examined at early (3 DPI), mid (6 DPI), and late disease (10 DPI, or the terminal time point in fatal cases). DPI, days post infection. Corresponding z-scores were plotted. Red indicates high expression; blue indicates low expression; white indicates no difference in expression; gray indicates insufficient transcripts mapping to the pathway.

Lastly, we measured serum IgG and IgM levels in vaccinated macaques to define the contribution of the humoral response to rVSV-N4CT1-MARV-GP-elicited protection. Only specifically vaccinated survivors formed substantial MARV-GP-specific IgG titers, with earlier detection of both immunoglobulin classes in subjects immunized 7 days prior to challenge (as early as the day of challenge in one subject) (**Fig 6A and 6B**). All surviving subjects had detectable IgG titers by 6 DPI, whereas no evidence of IgG was evident in vector control or non-surviving vaccinated macaques. Low IgM titers (1:100 to 1:800) generally declined during the convalescent stage conjointly with increasing moderate to high titers of IgG (1:400 to 1:12,800) (**Fig 6B**).

**Fig 6.**
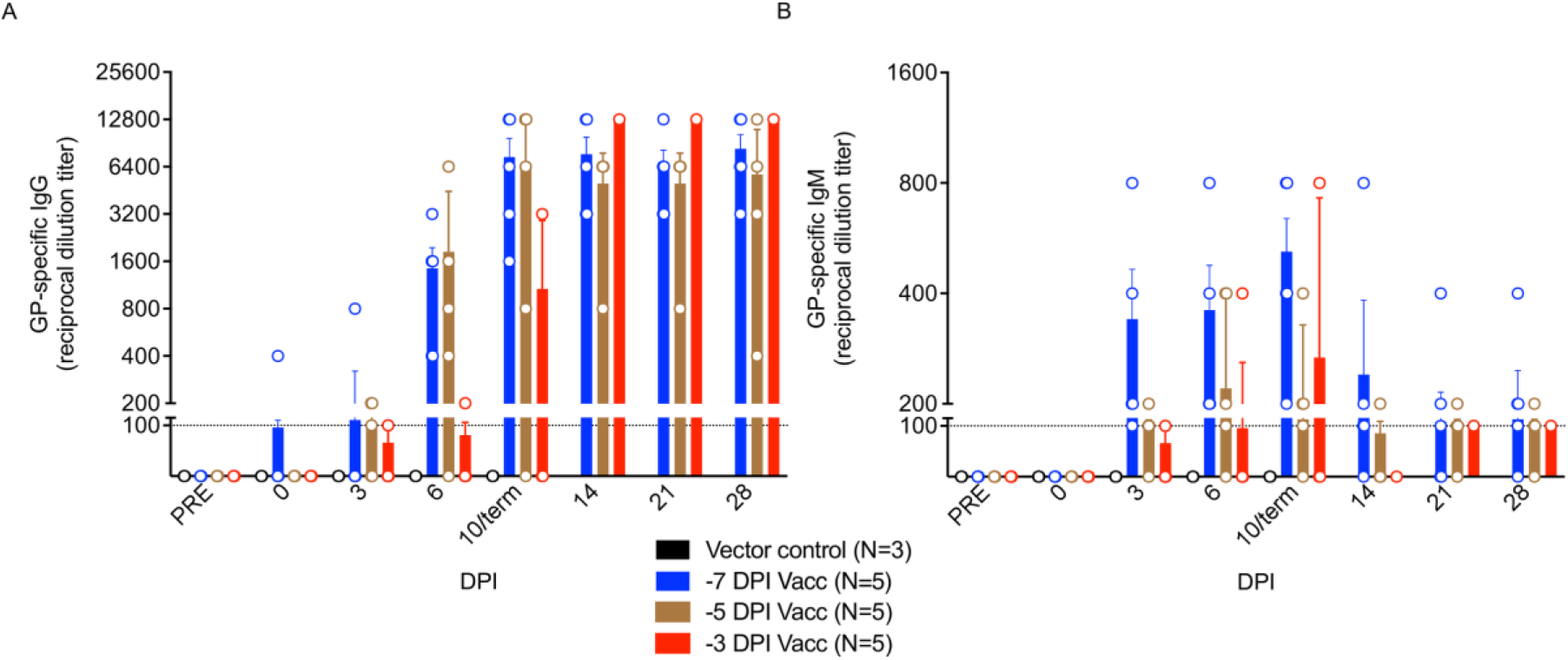
Reciprocal endpoint dilution titers of anti-MARV GP IgG and IgM in Vesiculovax-vaccinated cynomolgus macaques. MARV GP-specific **(A**) IgG and (**B**) IgM titers in vaccinated cohorts immunized with rVSV-N4CT1-MARV-GP at -7 DPI (blue; n=5), -5 DPI (brown; n=5), or -3DPI (red; n=5), or a vector control (black; n=3). The mean titer ± SEM is shown for each group at the denoted time points. Abbreviations: IgG, immunoglobulin G; IgM, immunoglobulin M; DPI, days post infection; term, terminal; MARV, Marburg virus; GP, glycoprotein; PRE, pre-vaccination baseline.

## Discussion

In a prior study, we showed that a highly attenuated rVSV-based vector, rVSV-N4CT1- MARV-GP, administered 20-30 minutes postexposure to a low dose MARV-Angola challenge afforded partial protection (60% survival) of NHPs [14]. As an extension of this previous work, we determined the minimum time needed between immunization and exposure to fully protect NHPs against MVD. Here, vaccination of macaques with a single 10 million PFU dose of rVSV-N4CT1-MARV-GP one week prior to MARV-Angola challenge resulted in lack of detectable viremia in these animals with protection from clinical disease and lethality. [27]. However, we detected changes in bloodwork of surviving subjects indicative of very mild disease in the absence of clinical signs. Immunization at 5 or 3 days before exposure resulted in 80% and 20% survival, respectively. Thus, this highly attenuated Vesiculovax vector provided comparable protection (100% survival) as the delta G vector as a rapid vaccine when administered one week before exposure. Although higher survival was reported for the delta G vector when given -3 days pre-challenge; no statistically significant difference was found between the two vaccines (log rank test, data not shown). Small group sizes used in the previous study make a direct comparison of vaccine efficacy difficult to interpret. Nevertheless, the fast-acting protection of rVSV-N4CT1-MARV-GP supports the use of Vesiculovax vaccines to combat future outbreaks of MVD. As a testament to this potential, ring vaccination with a similar EBOV-specific rVSV vector, Ervebo, reduced mortality and virus transmission during the 2013-2016 West Africa EBOV epidemic and 2018-2020 DRC EBOV outbreak [11, 12].

While the Ervebo “delta G (rVSVΔG)” vaccine appears highly efficacious, some undesirable vaccine-associated events have been reported in humans, particularly in individuals receiving high doses. Besides common vaccine reactions (injection site pain, fatigue, headache, muscle pain), these side effects included moderate to severe arthritis (with pain lasting a median of 8 days), postvaccination fever persisting for days, cutaneous vasculitis, and dermatitis manifesting as vesicular, maculopapular, or purpuric lesions distal to the inoculation site [28, 29]. Recovery of virus from synovial fluid and skin vesicles confirmed dissemination of the vaccine vector in peripheral tissues. Lowering the dose of Ervebo reduced reactogenicity, but also decreased immunogenicity and did not prevent arthritis, dermatitis, or cutaneous vasculitis in vaccinees. Although replication of the vaccine vector likely contributes to the rapid protection achieved with Ervebo, high viral loads throughout the body raises concerns about the potential short- and long-term effects of the vaccine. For these reasons, Ervebo is primarily indicated for reactive vaccination of individuals at high risk of exposure. A vaccine with comparably efficacy but a safer profile would be preferred for widespread use. Indeed, a rVSV-N4CT1 vector expressing EBOV GP protein similarly protected NHPs from lethal EBOV challenge despite a high degree of attenuation [41, 42] and was well tolerated and immunogenic in phase I clinical evaluation (https://clinicaltrials.gov/ct2/show/NCT02718469) [30]. The vaccine elicited robust EBOV GP-specific IgG responses and modest but balanced cellular immune responses. Importantly, no incidents of vaccine shedding, arthritis, nor skin rashes were reported in vaccinees at any dose tested including individuals given a high 18 million PFU dose. Based on these results, we anticipate rVSV-N4CT1-MARV-GP can similarly serve as a safe, immunogenic, and effective vaccine against MVD suitable for both preventive and reactive immunization.

The specific mechanisms of rapid rVSV-elicited protection are not fully elucidated but are thought to involve stimulation of both innate and adaptive immunity [13, 27, 43- 45]. Resistance to disease was strongly tied to early control of MARV replication. On the day of challenge, specifically vaccinated survivors compared to fatal cases tended to express higher levels of transcripts associated with interferon and RIG-I signaling, indicating a robust innate response may be needed to shape protective immunity. In survivors, innate signaling promptly resolved after MARV exposure unlike fatal cases concurrently with an accumulation of adaptive immunity-related transcripts, particularly in earlier vaccinated subjects. Administration of a non-specific rVSVN4CT1-HIV-gag1 vector control did not appear to delay the time-to-death and resulted in complete lethality of macaques following exposure to MARV-Angola, indicating MARV GP-specific responses modulate host resistance to MVD. This hypothesis is supported by the lack of substantial antibody titers and the absence of adaptive signaling in vector control subjects. Conversely, MARV GP-specific IgM and IgG titers were detected in all specifically vaccinated survivors by 0-6 DPI, and increased transcriptionally derived B cell, cytotoxic cell, and Th1 cell predicted frequencies were detected in these subjects as early as the day of challenge. Thus, humoral and cellular immune responses directed at the GP correlate with survival as we have previously shown [13].

Fatal outcome in vaccinated subjects corresponded with dramatic upregulation of transcripts encoding pro-inflammatory IP-10 and neutrophil-associated S100A8 and S100A9, which were previous reported to correlate with EBOV lethality in humans [36, 37] and in NHPs infected with *Bundibugyo ebolavirus* [33]. Similarly, lethality correlated with expression of immune checkpoint molecules as reported in other studies (*LAG3* [13], *CTLA4* [38]). Thus, certain factors associated with host susceptibility to MVD appear shared among various filovirus infections. The most significantly downregulated mRNA in vector control and vaccinated non-surviving subjects was complement factor D (*CFD*), a serine protease that cleaves complement factor B in the alternative complement pathway leading to downstream activation of the membrane attack complex [46]. As CFD is known to participate in opsonization and killing of pathogens, and viruses are commonly known to evade the complement system, this novel filovirus disease mechanism should be further explored.

Our transcriptomic results revealed NK cells and cytotoxic T cells are strongly implicated in rVSV-N4CT1-MARV-GP-associated fast-acting protection. Transcripts encoding cytoplasmic granules including perforin (*PRF1*), granzyme A (*GZMA*), granzyme B (*GZMA*), granzyme K (*GZMA*), and granzyme H (*GZMH*) were among the topmost upregulated molecules in vaccinated survivors. Not until late disease did we observe granzyme expression in fatal subjects. Activation of lymphocytes in non-survivors likely represents cytokine-mediated and not antigen-dependent T-cell activation as filovirus infection is known to interfere with antigen presentation [47, 48], and these subjects exhibited high viral loads. Increased expression of transcripts encoding killer cell lectin-type receptors (*KLRF1* (NKp80), *KLRB1* (CD161), *KLRD1* (CD94)) was also evident in surviving subjects. These receptors are ubiquitously expressed on NK cells and are involved in stimulation and regulation of cytotoxicity or self-nonself discrimination, further supporting this claim.

Previously, we showed higher overall NK cell frequencies and recruitment of a specific subset (CD16^+^) of NK cells were important for postexposure protection against MVD with rVSVΔG and rVSV-N2CT1 vectors [13]. CD16^+^ CD8a^+^ NK cells in NHPs represent the equivalent of the highly cytotoxic CD56^dim^ NK cell subset in humans [40]. These cells have limited cytokine-secreting potential but participate in killing virally infected cells or initiating antibody-dependent cell-mediated cytotoxicity (ADCC). Accordingly, depletion studies have revealed NK cell-intact mice survive longer after a mouse-adapted EBOV challenge, which is further enhanced by postexposure treatment with a rVSVΔG expressing EBOV GP (VSVΔG/EBOV GP) [49]. The authors of this study reported treatment with VSVΔG/EBOV GP treatment resulted in significantly higher NK cell-mediated cytotoxicity and IFN-γ secretion. Furthermore, recent research using a systems biology approach has identified NK cells as a primary correlate of antibody induction in humans receiving Ervebo. [44]. Specifically, the authors found that the frequency of CD56^bright^ NK cells on day 3 postvaccination and the expression of CXCR6 on CD56^dim^ NK cells on day 1 postvaccination positively correlated with antibody responses suggesting a potential role for antibody-mediated NK cell activation in vaccine-induced immune responses [50]. Thus, early NK cell differentiation status may dictate Fc-mediated activation of NK cells. This finding may be of relevance as antibody neutralizing capacity is not highly correlative of protection against MVD, and only low levels of neutralizing antibodies are typically detected following rVSV-MARV vaccination [13, 27].

Antibody neutralizing capacity and non-neutralizing antibody mechanisms were not assessed in this study, but future examinations should examine the role of ADCC and other Fc-mediated antibody effector functions in rVSV-N4CT1-MARV-GP-mediated protection against MVD.

Other immune components may also confer rVSV-N4CT1-MARV-GP protection. Results from clinical trials showed VSVΔG/EBOV GP elicited GP-specific CTLs, follicular T helper cells, and IFN-γ-secreting T helper cells [51–53]. Not surprisingly, Tbet (*TBX21*), the lineage-defining transcription factor for Th1 cells [54], was one of the most significantly upregulated transcripts in survivors 3-10 DPI in this study. Th1 cells secrete IFN-γ and IL-2 and represent a lineage of CD4^+^ effector T cells that promote cell-mediated immunity and defense against intracellular pathogens. As we’ve previously reported, postexposure survival against MVD following rVSV vaccination corresponded with increased polyfunctional IFN-γ^+^ and IL-2^+^ MARV GP-specific Th1 cells [13]. Other effector functions of Tbet include promotion of 1) immunoglobulin class switching, 2) the terminal differentiation of CD8+ T cells, 3) the maturation of NK cells, and 4) the secretion of cytoplasmic granules such as granzymes (46). Additionally, Tbet can protect the host from amplification of aberrant innate responses by dampening type-I IFN transcription factors and IFN-stimulated genes [55], which are highly expressed after MARV-Angola infection [56] and corresponded with lethal outcome in this study. Our pathway analysis supported activation of T cell responses and simultaneous repression of pathways involved in pattern recognition receptor and interferon signaling in line with this reasoning. Based on these collective findings, we infer virus-specific cytotoxic and helper T cells, along with plasma cells and NK cells, likely play a key role in defense against MVD.

While the molecular mechanisms involved in swift protection against MVD need additional clarification, we demonstrate that innate and adaptive immunity likely act in concert to elicit fast-acting protection. Further investigation is needed to determine how individual host responses to rVSV-N4CT1-MARV-GP vaccination alter susceptibility to MVD. In conclusion, the rapid protection mediated by rVSV-N4CT1-MARV-GP, along with promising safety and immunogenicity data from a phase I clinical evaluation with an analogous rVSV-N4CT1 vaccine, supports the testing of rVSV-N4CT1-MARV-GP in phase I clinical trials as the next step in developing an effective and safe vaccine against MARV. Our results indicate highly attenuated Vesiculovax vaccines may be suitable for ring vaccination during MVD outbreaks to save lives and reduce community transmission.

## Acknowledgements

The authors wish to thank the UTMB Animal Resource Center for husbandry support of laboratory animals. We wish to thank Drs. Kevin Melody and Abhishek Prasad for assistance with the animal experiments.

## Supporting Information Captions

**Fig S1.**
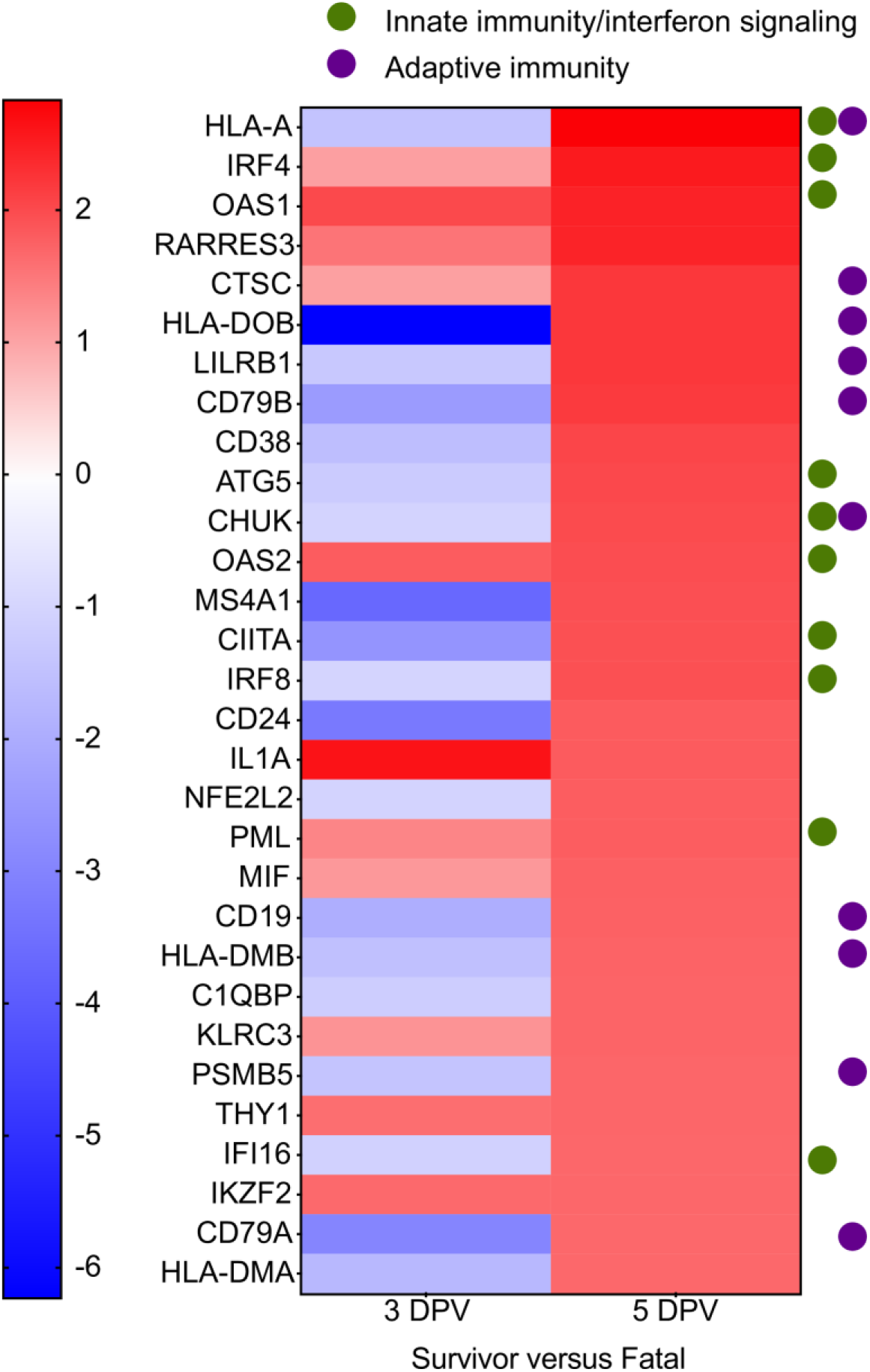
The topmost upregulated transcripts in Vesiculovax-vaccinated survivors on the day of challenge. Heatmap of the most upregulated mRNAs in survivor versus fatal subjects at five days post rVSV-N4CT1-MARV-GP vaccination (DPV) compared to 3 DPV. As only a single survivor exists in the 3 DPV group and a single fatal case exists in the 5 DPV group, statistical analysis was not performed. Red indicates high expression; blue indicates low expression; white indicates no difference in expression. Dots indicate whether specific mRNA is involved in adaptive immunity or innate immunity/interferon signaling.

**Fig S2.**
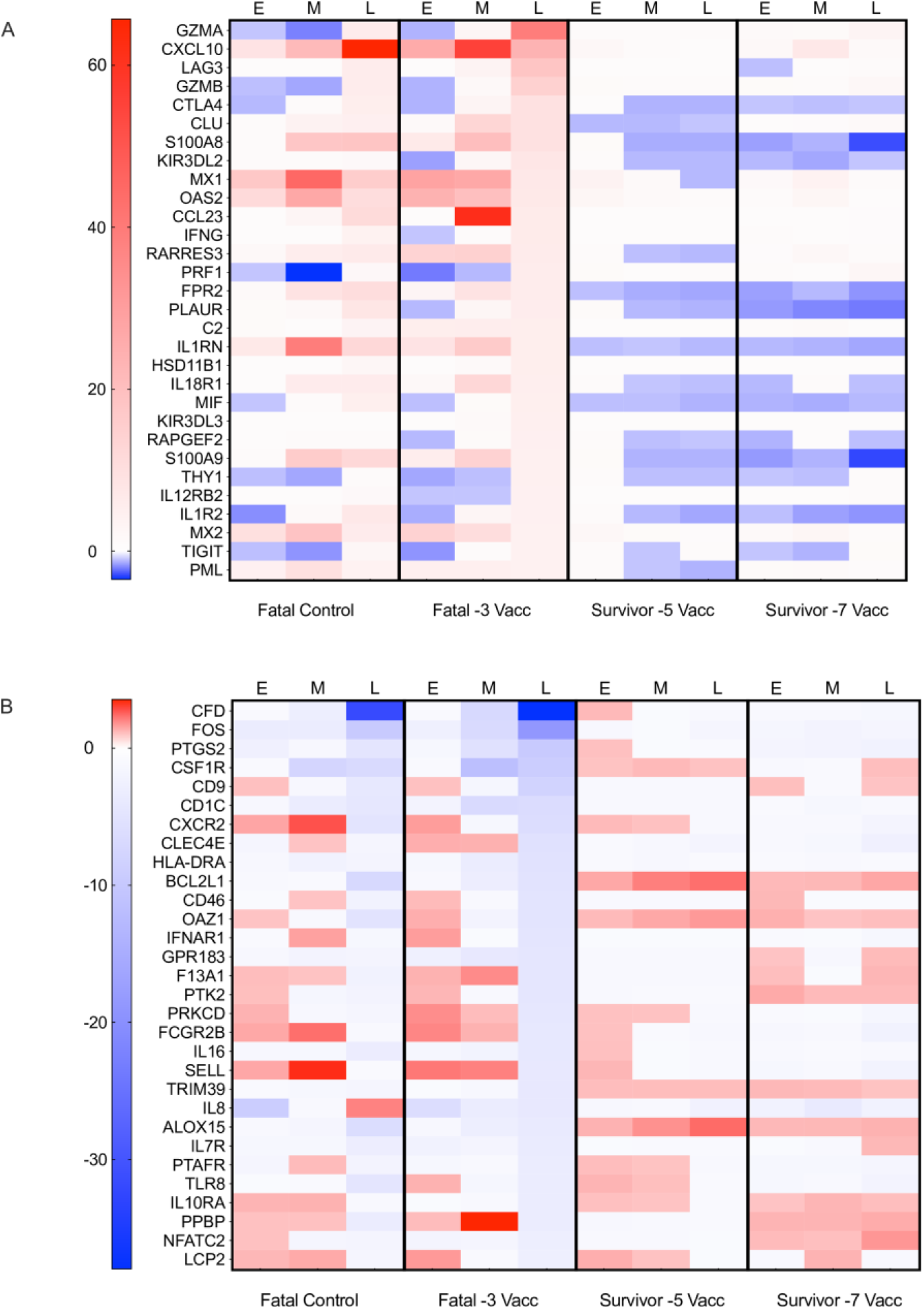
The topmost upregulated and downregulated transcripts in fatal subjects vaccinated with Vesiculovax vaccines. Heatmap of the most upregulated mRNAs in fatal subjects at five days post rVSV-N4CT1- MARV-GP vaccination (DPV) compared to 3 DPV. As only a single survivor exists in the 3 DPV group and a single fatal case exists in the 5 DPV group, statistical analysis was not performed. Red indicates high expression; blue indicates low expression; white indicates no difference in expression. Dots indicate whether specific mRNA is involved in adaptive immunity or innate immunity/interferon signaling. Abbreviations: Vacc, vaccinated; E, early disease (3 DPI); M, mid disease (6 DPI); L, late disease (10 DPI or terminal timepoint).

**Fig S3.**
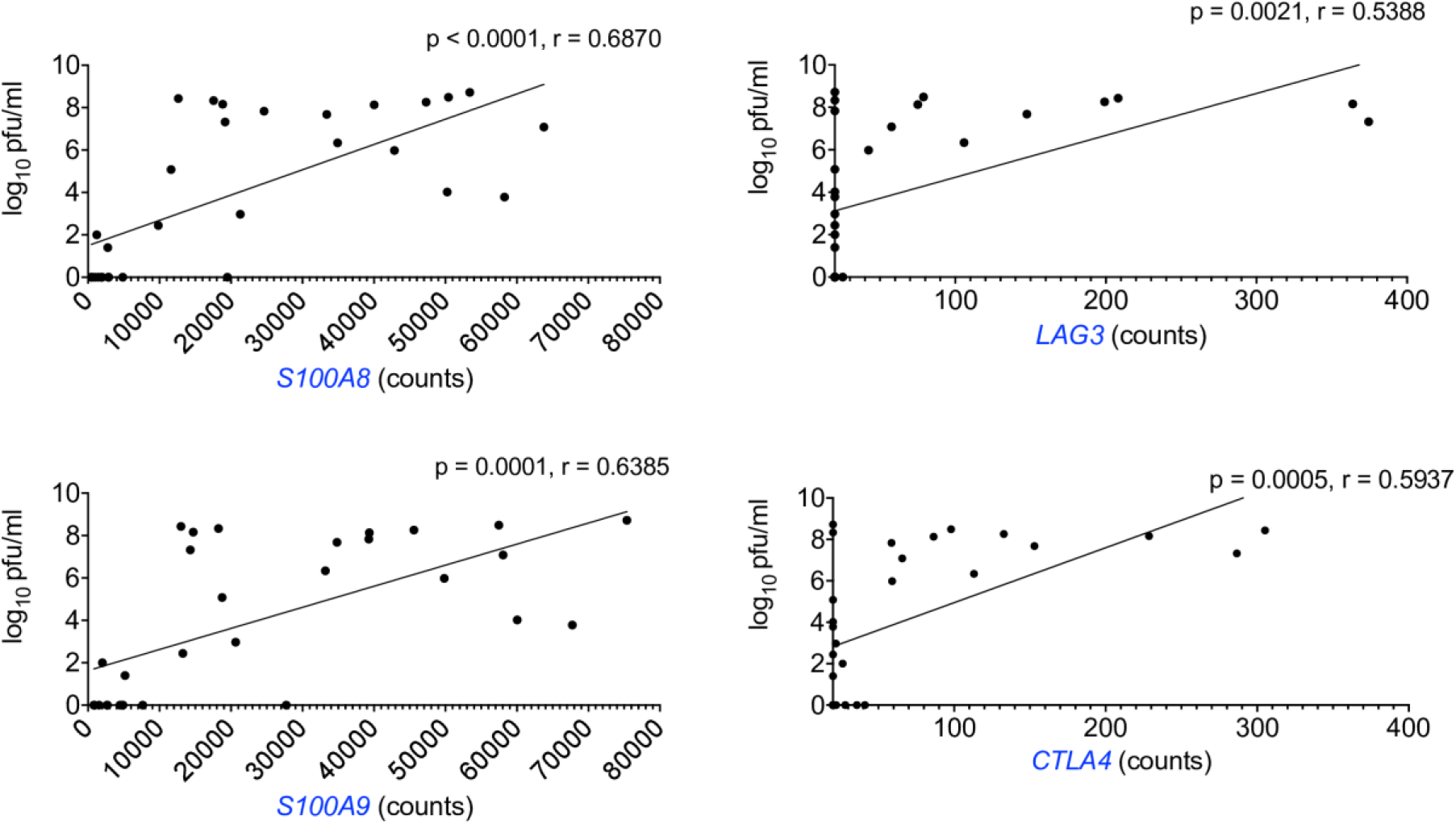
Transcriptional correlates associated with lethal outcome in Vesiculovax- vaccinated macaques exposed to MARV-Angola. Pearson correlation plots for calprotectin (S100A8 and S100A9) and immune checkpoint (LAG3 and CTLA4) transcripts (Benjamini-Hochberg false discovery rate (FDR) corrected p-value less than 0.05).

**S1 Table.**
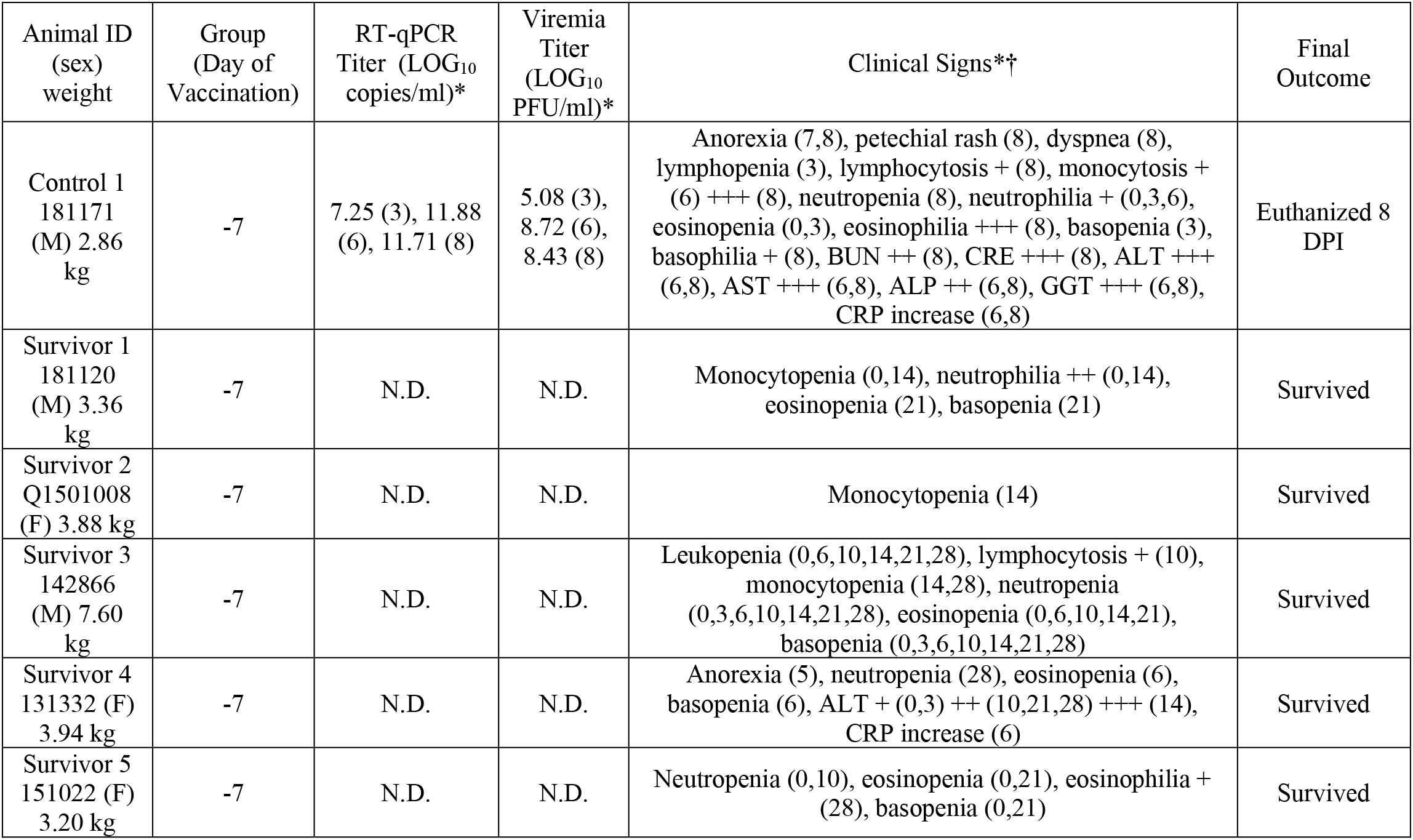
Clinical findings in MARV-exposed cynomolgus macaques immunized with Vesiculovax vaccine 7 days prior to challenge. Macaques were immunized with a vector control (black; n=1) or rVSV-N4CT1-MARV- GP vaccine at -7 DPI (blue; n=5). *Day after MARV challenge is in parentheses up to the 28 DPI study endpoint. †Fever is defined as a temperature greater than 2.5 °F above baseline, at least 1.5 °F above baseline and ≥ 103.5 °F, or 1.1 °F above baseline and ≥ 104°F. Leukopenia, thrombocytopenia, and lymphopenia are defined by a > 40% drop in numbers of leukocytes, platelets, and lymphocytes, respectively. Leukocytosis, monocytosis, and granulocytosis are defined as a ≥ two-fold increase in leukocytes, monocytes, and granulocytes, respectively. Crosses indicate increases in liver enzymes (ALT, AST, ALP, GGT) or renal function test values (BUN, CRE): 2- to 3-fold increase, +; >3- up to 5-fold increase, ++; >5-fold increase, +++. Abbreviations: M, male; F, female; kg, kilogram; PFU, plaque-forming units; MARV, Marburg virus; BUN, blood urea nitrogen; CRE, creatinine; ALT, alanine aminotransferase; AST, aspartate aminotransferase; ALP, alkaline phosphatase; GGT, gamma-glutamyltransferase; CRP, c- reactive protein; DPI, days post infection.

| Animal ID<br>(sex)<br>weight | Group<br>(Day of<br>Vaccination) | RT-qPCR<br>Titer (LOG<sub>10</sub><br>copies/ml)* | Viremia<br>Titer<br>(LOG<sub>10</sub><br>PFU/ml)* | Clinical Signs*† | Final<br>Outcome |
| --- | --- | --- | --- | --- | --- |
| Control 1<br>181171<br>(M) 2.86<br>kg | -7 | 7.25 (3), 11.88<br>(6), 11.71 (8) | 5.08 (3),<br>8.72 (6),<br>8.43 (8) | Anorexia (7,8), petechial rash (8), dyspnea (8), lymphopenia (3), lymphocytosis + (8), monocytosis + (6) +++ (8), neutropenia (8), neutrophilia + (0,3,6), eosinopenia (0,3), eosinophilia +++ (8), basopenia (3), basophilia + (8), BUN ++ (8), CRE +++ (8), ALT +++ (6,8), AST +++ (6,8), ALP ++ (6,8), GGT +++ (6,8), CRP increase (6,8) | Euthanized 8<br>DPI |
| Survivor 1<br>181120<br>(M) 3.36<br>kg | -7 | N.D. | N.D. | Monocytopenia (0,14), neutrophilia ++ (0,14), eosinopenia (21), basopenia (21) | Survived |
| Survivor 2<br>Q1501008<br>(F) 3.88 kg | -7 | N.D. | N.D. | Monocytopenia (14) | Survived |
| Survivor 3<br>142866<br>(M) 7.60<br>kg | -7 | N.D. | N.D. | Leukopenia (0,6,10,14,21,28), lymphocytosis + (10), monocytopenia (14,28), neutropenia (0,3,6,10,14,21,28), eosinopenia (0,6,10,14,21), basopenia (0,3,6,10,14,21,28) | Survived |
| Survivor 4<br>131332 (F)<br>3.94 kg | -7 | N.D. | N.D. | Anorexia (5), neutropenia (28), eosinopenia (6), basopenia (6), ALT + (0,3) ++ (10,21,28) +++ (14), CRP increase (6) | Survived |
| Survivor 5<br>151022 (F)<br>3.20 kg | -7 | N.D. | N.D. | Neutropenia (0,10), eosinopenia (0,21), eosinophilia + (28), basopenia (0,21) | Survived |

**S2 Table.**
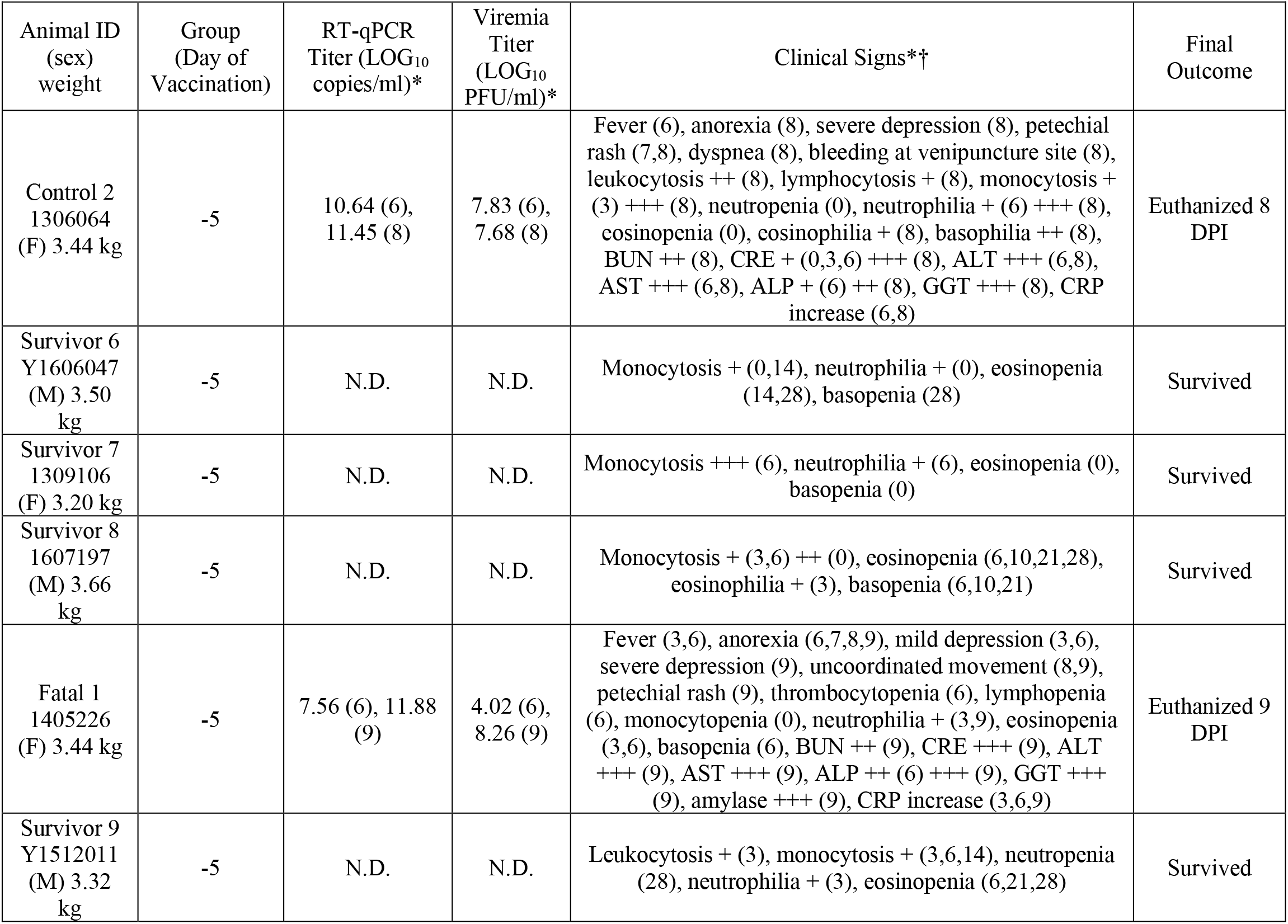
Clinical findings in MARV-exposed cynomolgus macaques immunized with Vesiculovax vaccine 5 days prior to challenge. Macaques were immunized with a vector control (black; n=1) or rVSV-N4CT1-MARV- GP vaccine at -5 DPI (brown; n=5). *Day after MARV challenge is in parentheses up to the 28 DPI study endpoint. †Fever is defined as a temperature greater than 2.5 °F above baseline, at least 1.5 °F above baseline and ≥ 103.5 °F, or 1.1 °F above baseline and ≥ 104°F. Leukopenia, thrombocytopenia, and lymphopenia are defined by a > 40% drop in numbers of leukocytes, platelets, and lymphocytes, respectively. Leukocytosis, monocytosis, and granulocytosis are defined as a ≥ two-fold increase in leukocytes, monocytes, and granulocytes, respectively. Crosses indicate increases in liver enzymes (ALT, AST, ALP, GGT) or renal function test values (BUN, CRE): 2- to 3-fold increase, +; >3- up to 5-fold increase, ++; >5-fold increase, +++. Abbreviations: M, male; F, female; kg, kilogram; PFU, plaque-forming units; MARV, Marburg virus; BUN, blood urea nitrogen; CRE, creatinine; ALT, alanine aminotransferase; AST, aspartate aminotransferase; ALP, alkaline phosphatase; GGT, gamma-glutamyltransferase; CRP, c- reactive protein; DPI, days post infection.

**S3 Table.**
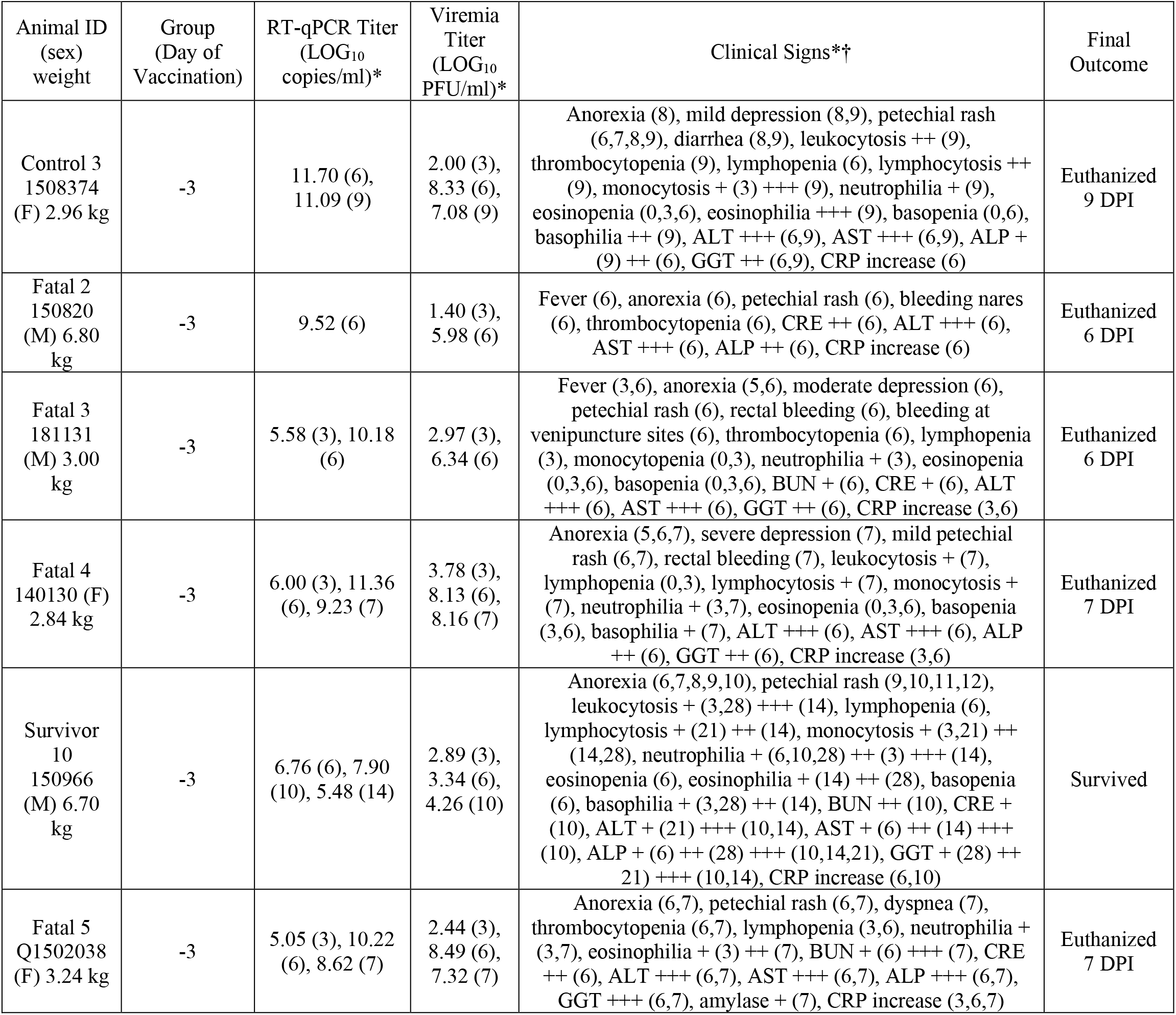
Clinical findings in MARV-exposed cynomolgus macaques immunized with Vesiculovax vaccine 3 days prior to challenge. Macaques were immunized with a vector control (black; n=1) or rVSV-N4CT1-MARV- GP vaccine at -3DPI (red; n=5). *Day after MARV challenge is in parentheses up to the 28 DPI study endpoint. †Fever is defined as a temperature greater than 2.5 °F above baseline, at least 1.5 °F above baseline and ≥ 103.5 °F, or 1.1 °F above baseline and ≥ 104°F. Leukopenia, thrombocytopenia, and lymphopenia are defined by a > 40% drop in numbers of leukocytes, platelets, and lymphocytes, respectively. Leukocytosis, monocytosis, and granulocytosis are defined as a ≥ two-fold increase in leukocytes, monocytes, and granulocytes, respectively. Crosses indicate increases in liver enzymes (ALT, AST, ALP, GGT) or renal function test values (BUN, CRE): 2- to 3-fold increase, +; >3- up to 5-fold increase, ++; >5-fold increase, +++. Abbreviations: M, male; F, female; kg, kilogram; PFU, plaque-forming units; MARV, Marburg virus; BUN, blood urea nitrogen; CRE, creatinine; ALT, alanine aminotransferase; AST, aspartate aminotransferase; ALP, alkaline phosphatase; GGT, gamma-glutamyltransferase; CRP, c-reactive protein; DPI, days post infection.

## Availability of data and materials

The datasets used and/or analyzed during the current study are available from the corresponding author upon request.

## Author contributions

TWG, MAE, JHE, and DM conceived and designed the study. CG, TEL, and DM designed the vaccine vectors and did preparative work. DJD, JBG, and TWG performed the challenge experiments. CW, RWC, DJD, JBG, and TWG performed animal procedures and clinical observations. VB and KNA performed the clinical pathology assays. VB performed the plaque assays. KNA performed the PCR assays. CW performed the transcriptomic assays and bioinformatics. CW performed the ELISAs. KAF performed the necropsies and gross pathology analysis. All authors analyzed the data. CW wrote the paper. RWC, TWG, and DM edited the paper. All authors had access to the data and approved the final version of the manuscript.

